# Avid binding by B cells to the *Plasmodium* circumsporozoite protein repeat suppresses responses to protective subdominant epitopes

**DOI:** 10.1101/2020.01.12.903682

**Authors:** Fiona J. Lewis, Deepyan Chatterjee, Joe Kaczmarski, Xin Gao, Yeping Cai, Hayley A. McNamara, Henry J. Sutton, Colin J. Jackson, Ian A. Cockburn

## Abstract

Antibodies targeting the NANP/NVDP repeat domain of the *Plasmodium falciparum* circumsporozoite protein (CSP_Repeat_) can confer protection against malaria. However, it has also been suggested that this repeat domain exists as a decoy that distracts the immune system from mounting protective responses targeting other domains of CSP. Here we show that B cell responses to the repeat domain are indeed ∼10 fold higher than responses to the N- and C-terminal regions of CSP after sporozoite immunization. We investigated the role of the number of CSP_Repeat_-specific naïve precursor B cells and high avidity binding by B cells in driving the immunodominance of the CSP_Repeat_. Using adoptive transfer of germline precursors specific for the CSP_Repeat_, we found that increasing precursor number did indeed increase the responses to the repeat region, but not to the detriment of responses to other epitopes. To investigate the role of avid binding by B cells to the CSP_Repeat_ in driving immunodominance we generated CSP9: a truncated CSP molecule with just 9 NANP repeats. Compared to near full length CSP molecules, CSP9 induced lower BCR signalling in CSP_Repeat_-specific cells and induced stronger responses to non-repeat epitopes. Finally, we found mice immunized with CSP9 molecules were strongly protected against mosquito bite challenge. Collectively these data demonstrate that the CSP_Repeat_ does function as an immunodominant decoy and that truncated CSP molecules may be a promising avenue for future malaria vaccines.

**Significance Statement:** Malaria kills approximately 420,000 individuals each year(1). Our best vaccine, RTS,S/AS01 is based on the circumsporozoite protein that coasts the surface of the parasite. However, this vaccine is only partially protective. Here we show that responses to a repeat region in the circumsporozoite dominate the immune response. However, immunizing with a circumsporozoite protein with a shortened repeat region induces a more diverse immune response, which could be an avenue to make better malaria vaccines.

## Introduction

Our most advanced malaria vaccine RTS,S/AS01 aims to induce antibodies that target the repeat region of the circumsporozoite protein (CSP), which covers the surface of the *Plasmodium* sporozoite (2-4). The rationale for this approach comes from the observation that immunization with radiation attenuated sporozoites confers sterile protection against malaria, and that the humoral response induced by irradiated sporozoites is dominated by anti-CSP antibodies (5-8). Early studies demonstrated that monoclonal antibodies (mAbs) targeting the repeat regions of the CSP molecule (CSP_Repeat_) protected mice against challenge with the rodent parasite *P. berghei* (9-11). More recently, human monoclonal antibodies targeting the *P. falciparum* CSP_Repeat_ have also been shown to be protective in preclinical mouse models (12-14).

Despite the protective capacity of CSP_Repeat_ specific Abs, it has also been argued that the CSP_Repeat_ is an immunodominant “decoy” distracting the immune system from making protective responses against other epitopes within CSP or other proteins on the sporozoite surface (15, 16). Evidence for the immunodominance of the CSP_Repeat_ initially came from early studies which showed that a short (NANP)_3_ peptide based on this domain could absorb most sporozoite binding activity of sera from hyperimmune individuals (8). In support of the concept that the responses to CSP_Repeat_ are sub-optimal, large amounts of anti-CSP_Repeat_ mAbs are required for protection in preclinical challenge models, while in RTS,S clinical trials protection requires very high amounts (>50 μg/ml) of anti-CSP_Repeat_ antibody (12-14, 17). In contrast, antibody responses to other regions of CSP are less well understood. One small epidemiological study associated increased levels of antibodies targeting a truncated CSP_Nterm_ peptide with protection from clinical disease (18). Subsequently, a mouse mAb, 5D5, targeting an epitope within the N terminus of CSP was found to be protective against sporozoite challenge (19). More recently, human mAbs targeting the junction between the CSP_Nterm_ and CSP_Repeat_ were found to be protective (12, 13). Antibodies targeting the C-terminal domain CSP (CSP_Cterm_) have been associated with protection by the RTS,S vaccine in clinical trials (20, 21), though individual mAbs targeting this domain have not been found to confer protection (14).

There has been increased interest in the factors that drive B cell immunodominance and how these can be manipulated for improved vaccination outcomes. Recent findings in influenza and HIV immunology have revealed the existence of broadly neutralising antibodies (bnAbs), however these target rare subdominant epitopes. In HIV it was recently shown that transferred B cells carrying a germline version of the bnAb VRC01 could be induced to compete successfully in germinal centers (GCs) if the number of naïve precursors was artificially increased or if stimulated with a high polyvalent antigen that bound the B cells with greater avidity (22). For influenza it has been shown that broadly neutralizing responses to the stem regions of haemagglutinin (HA) can be favored over responses to the immunodominant - but highly variable - head region by immunization with stem-only constructs, even if delivered alongside full length HA (23).

Given the highlighted roles for antigen polyvalency and precursor numbers in driving B cell immunodominance, we investigated whether these factors drive the CSP_Repeat_ to be immunodominant. The finding that CSP_Repeat_ can be bound avidly by B cells from a range of immunoglobulin gene families suggested that there may be high numbers of precursors for this domain (12, 13, 24, 25). More suggestively, we and others have shown that the CSP repeat can be bound by 6 or more specific antibodies (12, 24, 26, 27), and it has been demonstrated that the repeat can crosslink multiple BCRs to enhance B cell signalling (28). Accordingly, we tested the roles of these factors in driving the dominance of the Ab response against the CSP_Repeat_ and determined if we could manipulate the immunodominance hierarchy to develop better vaccination protocols.

## Results

### The circumsporozoite protein repeat domain is immunodominant

To formally test the immunodominance of responses to the CSP_Repeat_ over the CSP_Nterm_ and CSP_Cterm_ we immunized mice with irradiated Pb-PfSPZ parasites which carry a full length (4NVDP/38NANP) *P. falciparum* CSP gene in place of the endogenous *P. berghei* CSP (Fig. S1) (29). At days 4, 7, 14 and 28 post-immunization sera were taken for antibody analysis by ELISA with domain-specific peptides, and spleens were taken for cellular analysis by flow cytometry. IgG responses to the CSP_Repeat_ were significantly higher than responses to either of the other domains, with a significant response developing to the CSP_Cterm_ only after 28 days (Fig. 1A).

**Fig. 1.**
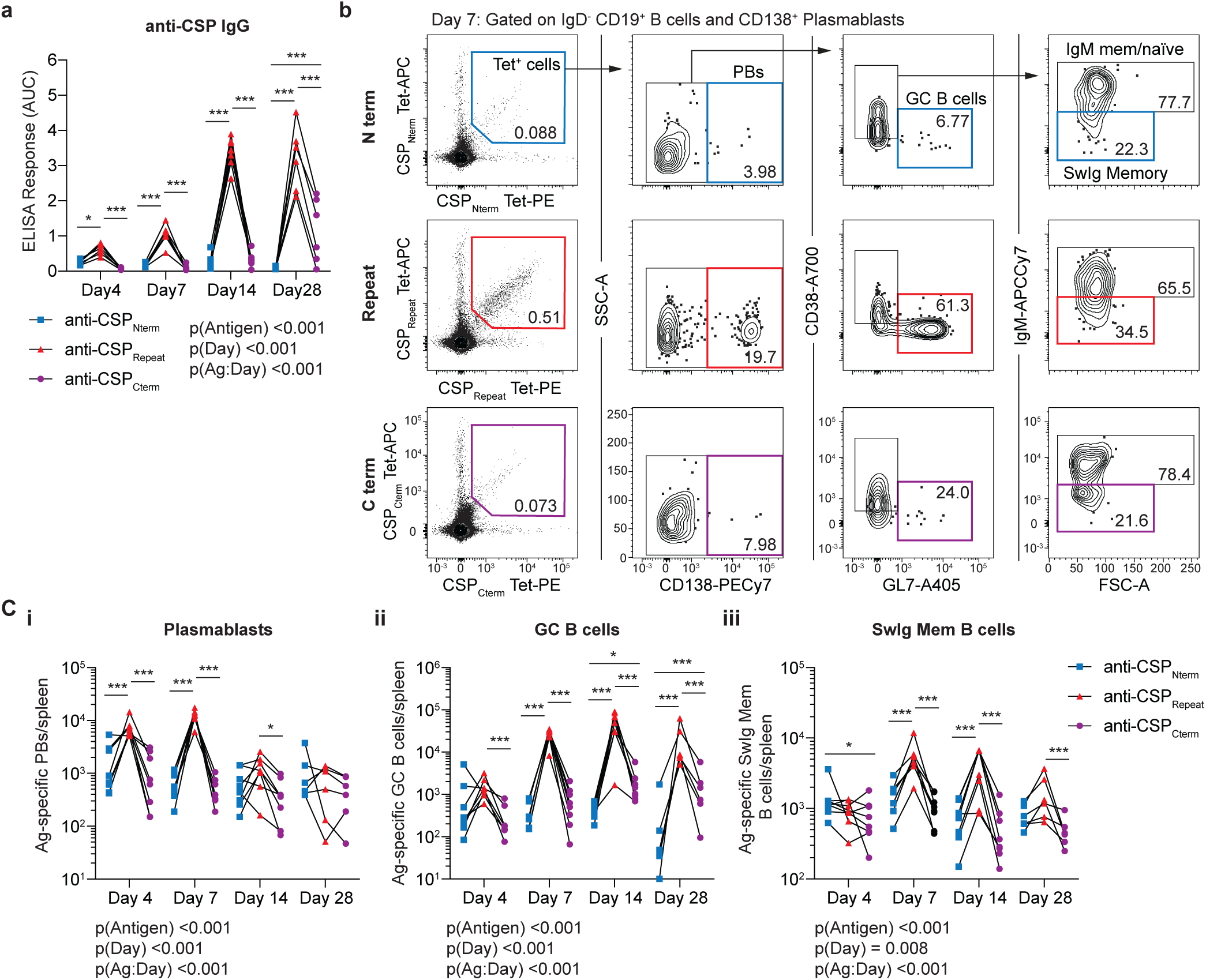
Responses to the CSP_Repeat_ are immunodominant over responses to other domains. C57BL/6 mice were immunized with 5 x10^4^ irradiated Pb-PfSPZ sporozoites, blood and spleens were taken at 4, 7, 14 and 28 days post-immunization for analysis by ELISA and flow cytometry using probes specific for each domain of CSP (CSP_Cterm_, CSP_Repeat_ and CSP_Nterm_). (A) IgG responses to each domain measured by ELISA, data shown as area under the curve. (B) Representative flow cytometry plots from a single mouse at the day 7 timepoint showing the gating of PBs, GC B cells and SwIg Mem for antigen specific IgD-B cells identified using tetramers specific for the CSP_Cterm_, CSP_Repeat_ and CSP_Nterm_; values are percentages (C) Quantifications of the absolute numbers of IgD- B cells using NANP, R1+ and N_81-91_ antibody panels. (C) Absolute numbers of (i) Plasmablasts, (ii) GC B cells and (iii) SwIg memory cells in each mouse for each antigen (domain). Data for all panels are represented as mean ± SD pooled from two independent experiments (n=3-5 mice/timepoint/experiment); all data were analyzed via 2-way ANOVA, with experiment and mouse included in the model as fixed factors, ANOVA p values are listed below or adjacent to each graph. Pairwise comparisons were performed using a Tukey post-test and significant pairwise comparisons are represented as symbols; * p<0.05, ** p<0.01, *** p<0.001.

One concern is that ELISA measurements of antibody responses to the different domains are not directly comparable. Therefore, we used tetramer probes to track the total numbers of B cells responding to each domain of CSP over time, and their phenotype by flow cytometry (Fig. 1B, Fig. S2A). The response to sporozoites is characterized by an early plasmablast (PB) response that wanes and leaves a prolonged GC reaction (24). Four days after immunization of mice with Pb-PfSPZ, the number of CSP_Repeat_^+^ CD138^+^ PBs was ∼10 fold higher than the number of CSP_Nterm_^+^ or CSP_Cterm_^+^ PBs (Fig. 1Ci). By day 7, a pronounced GC reaction developed and the number of CSP_Repeat_^+^ GL7^+^ B cells was ∼10 fold higher than responses to the other domains, which was sustained until day 28 (Fig 1Cii). We further analyzed the number of IgD^-^ IgM^-^ (Switched Ig; SwIg) CD38^+^ memory B cells and found that the immunodominance of the response to the CSP_Repeat_ response extended into the memory phase (Fig 1Ciii).

### Increasing anti-CSP precursor B cell number does not supress responses to other antigens

A previous study highlighted the fact that increasing the number of precursors for an antigen led to a concomitant increase in the number of cells entering GCs; the same study also highlighted roles for antigen valency in allowing responses to successfully compete in the GC (22). To investigate the roles of precursor number and antigen valency we developed a system in which we could modulate the number of precursors or the valency of two competing antigens in the context of CSP. We achieved this by conjugating the 4-hydroxy-3-nitrophenyl (NP) acetyl-hapten to recombinant CSP via crosslinking on lysine. Since lysine residues are not found in the CSP repeat region, the NP-hapten would bind exclusively to the N and C terminal domains (Fig. S3A). For these experiments we used a slightly truncated CSP molecule carrying 27 repeats (3NVDP and 24 NANP), designated CSP27 (Fig S1). As proof of principle we found that immunization with CSP27 conjugated to 2 NP moieties (CSP27-NP2) was able to induce strong anti-NP and anti-CSP_Repeat_ IgG responses (Fig. S3). We were then able to modulate the number of precursors for CSP_Repeat_-specific B cells using Ig knockin Igh^g2A10^ B cells which carry the germline heavy chain of the CSP_Repeat_ specific 2A10 antibody (McNamara *et al.* submitted). Similarly, we could modulate the number of NP-specific B cells using the established B1-8^hi^ mouse system (30, 31). Finally, we could alter the valency of the response to NP or the repeat by conjugating more NP molecules per CSP, or by reducing the length of the CSP_Repeat_ domain.

To determine the role of precursor number in driving immunodominance we adoptively transferred defined numbers of CSP_Repeat_ tetramer^+^, CD45.1 Igh^g2A10^ cells into CD45.2 C57BL/6 mice (Fig. S2B). Mice were then immunized with CSP27-NP2, IgG responses were measured 7,14 and 21 days post immunization, and the number of NP and CSP_Repeat_ specific cells were quantified by flow cytometry on day 21 (Fig. 2A). We hypothesized that increased anti-CSP_Repeat_ precursor number would not only increase the response to the CSP_Repeat_ but also suppress the NP immune response. As expected, increasing the number of CSP_Repeat_-specific precursors increased the total number of CSP_Repeat_ binding B cells and CSP_Repeat_-specific GC B cells responding to this antigen (Fig. 2B and C). Perhaps surprisingly, the magnitude of the antibody response was unaltered (Fig. 2D). However, there was no concomitant decrease in the overall B cells and GC B cell response to NP (Fig. 2B-C), and the magnitude of the IgG antibody response to NP was also unaffected by the addition of Igh^g2A10^ cells (Fig. 2E). Finally, because we could distinguish our transferred cells from the endogenous response by the expression of CD45.1, we were able to determine that the endogenous response to CSP was not supressed by the addition of enhanced number of germline precursors specific for CSP (Fig 2F).

**Fig. 2.**
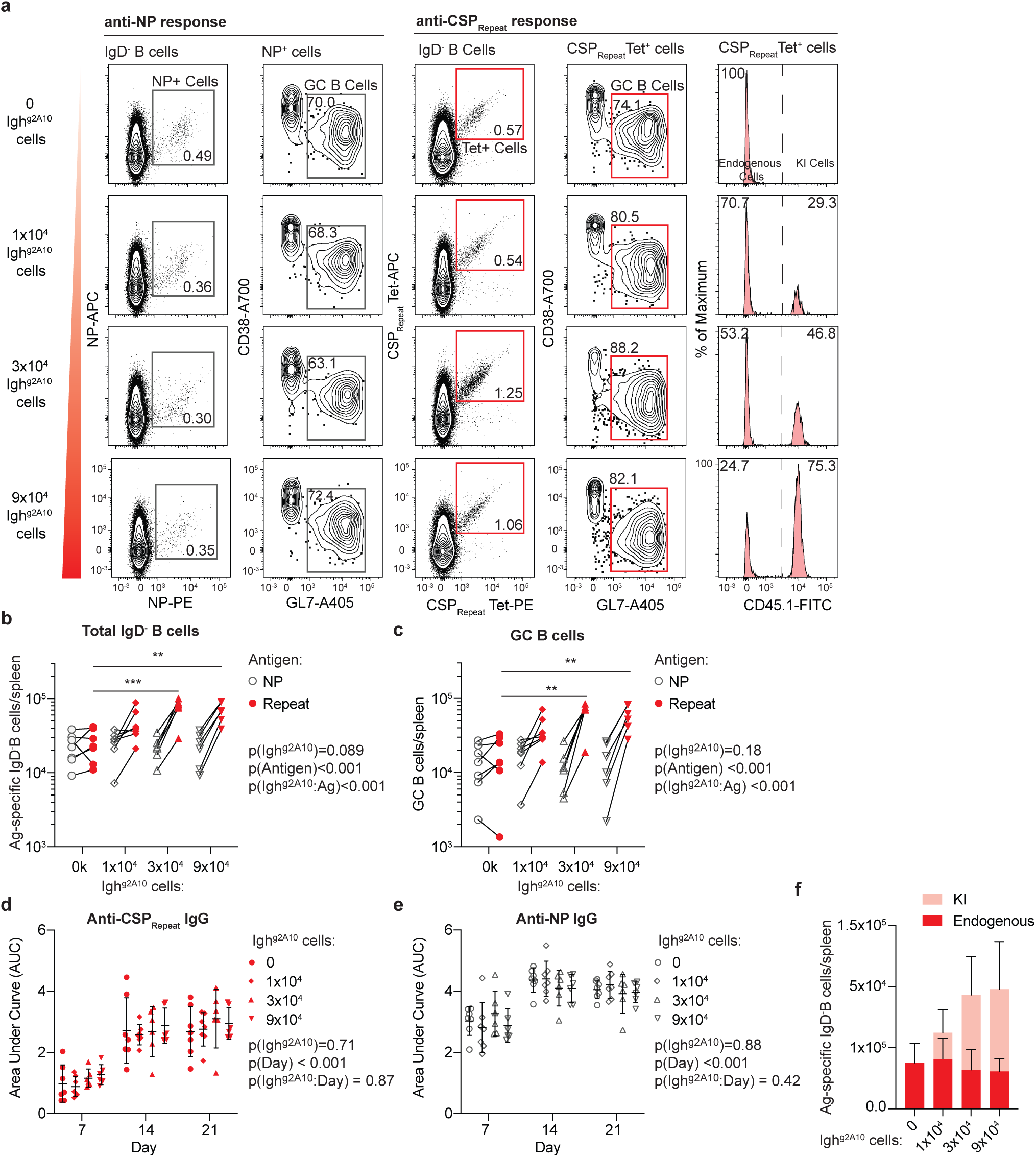
Increasing CSP_Repeat_-specific precursor number does not alter immunodominance. 0, 1×10^4^, 3×10^4^, or 9×10^4^ of CD45.1 Igh^g2A10^ cells were adoptively transferred into C57Bl/6 mice followed by immunization with 30 µg CSP27-NP2 in alum. Sera were taken on days 7, 14 and 21 and spleens analyzed 21 days post-immunization. (A) Representative flow cytometry plots showing gating of total IgD^-^ and GC B cells specific for NP or the CSP_Repeat_; values are percentages (B) Absolute numbers of NP probe^+^ and CSP_Repeat_ tetramer^+^ IgD^-^ B cells. (C) Absolute numbers of NP probe^+^ and CSP_Repeat_ tetramer^+^ GC B cells. (D) Total IgG response to CSP_Repeat_ measured via (NANP)_9_ ELISA. (E) Total IgG response to NP measured via NP(14)BSA ELISA. (F) Absolute numbers of CSP_Repeat_ tetramer^+^ CD45.1^+^ Igh^g2A10^ and CD45.1^-^ endogenous cells. Data are represented as mean ± SD pooled from two independent experiments (n≥3 mice/group/experiment); all data were analyzed via 2-way ANOVA, with experiment and mouse included in the model as fixed factors. ANOVA p values are listed below or adjacent to each graph. Pairwise comparisons were performed using a Tukey post-test and significant values are represented as symbols; * p<0.05, ** p<0.01, *** p<0.001.

We also performed the converse experiment and altered the number of anti-NP precursors relative to the number of CSP_Repeat_ specific cells via the addition of B1-8^hi^ cells (Fig. S2C). Notably, B1-8^hi^ cells differ from Igh^g2A10^ cells not only in their specificity but also because they carry the high affinity mature *Ighv1-72* heavy chain that confers strong binding to NP. Nonetheless, in agreement with the previous experiment, antibody titres to the CSP_Repeat_ were generally unaffected by the transfer of additional naive B-18^hi^ cells (Fig S4A), while increasing the number of NP cells had no significant effect on the number of CSP_Repeat_ specific B cells responding and becoming GC B cells (Figure S5B-D). However, in contrast to the previous experiment, the additional of B1-8^hi^ cells did not increase the overall magnitude of the antigen specific B cell and GC response to NP itself; rather, the transferred high affinity B1-8^hi^ B cells displaced the endogenous cells from the response though this effect rapidly saturated after the transfer of just 1×10^4^ B1-8^hi^ cells (Fig S4E). Again, in contrast to the previous experiment, this displacement resulted in higher titres of antibodies targeting NP overall, perhaps due to the high affinity of the transferred cells (Fig. S4F).

### Reducing the valency of immunodominant antigens allows subdominant responses to expand

Given the repeating nature of the CSP_Repeat_ we next tested whether the ability of long CSP molecules to crosslink multiple B cell receptors (BCRs) might drive the immunodominance of this domain. Accordingly, we developed a construct that carried just 9 NANP repeats (CSP9; Fig. S1). Using a surface plasmon resonance saturation experiment, we found that CSP9 could only bind 2-3 2A10 antibodies, compared to the 5-6 bound by CSP27, which is in line with previous structural and biophysical data from our laboratory (Fig. 3A). Importantly, this reduced binding corresponded to a reduction in BCR signalling as calcium fluxes were lower when Igh^g2A10^ cells were pulsed with CSP9 compared to CSP27 (Fig. 3B-C).

**Fig. 3.**
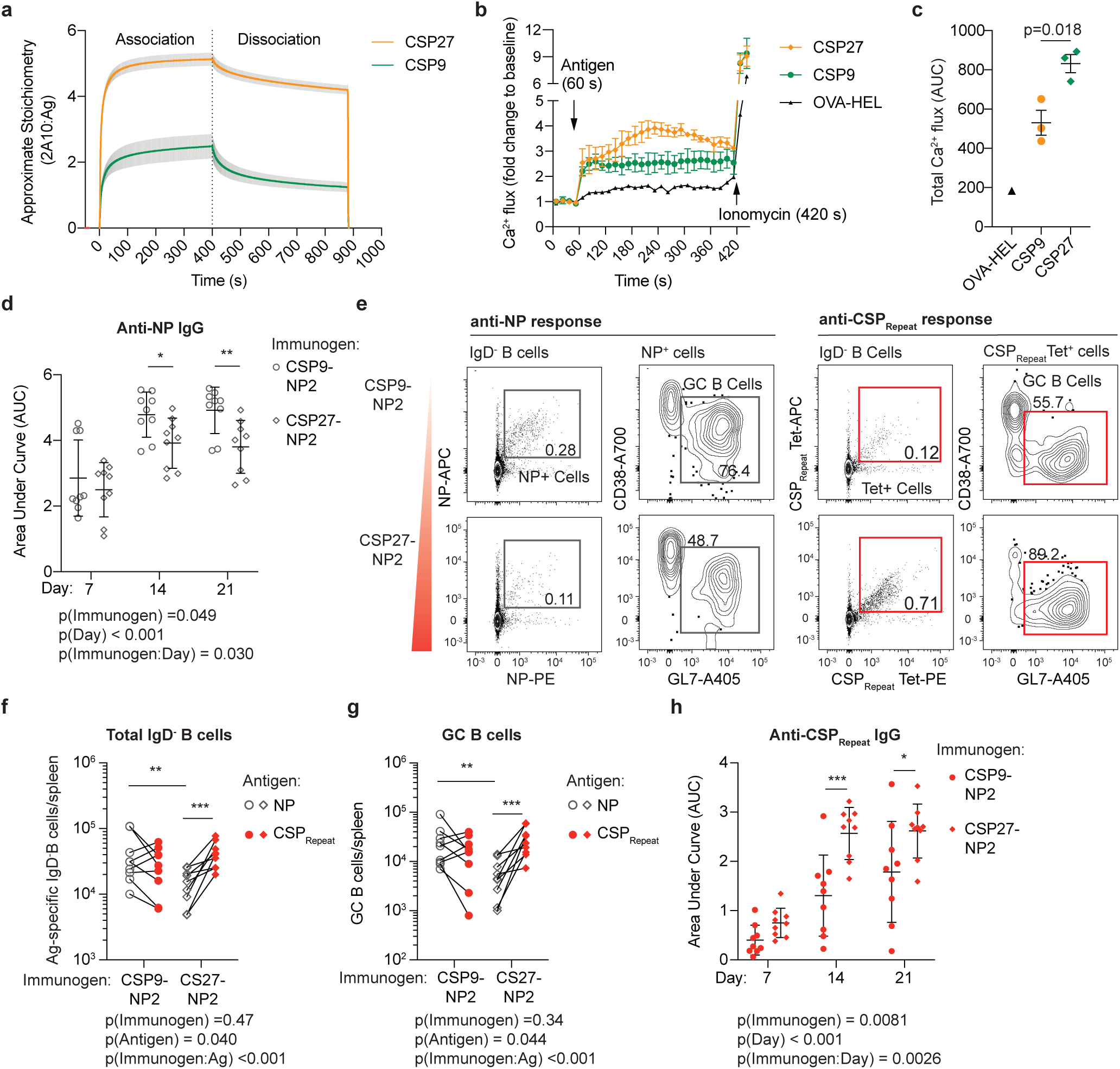
Decreasing the valency of the CSP_Repeat_ alters the immunodominance hierarchy. Recombinant CSP9 was purified and conjugated to NP at a 1:2 ratio to generate CSP9-NP2, mice were immunized with either CSP9-NP2 (23 µg) or CSP27-NP2 (30 µg) in alum; sera were taken on days 7, 14 and 21 and spleens analyzed 21 days post-immunization. (A) Approximate binding stoichiometry of the 2A10:Ag complex formed when a saturating concentration (2 μM) of mAb 2A10 was passed over immobilized CSP27 or CSP9; data shows mean ± SD of 2 technical replicates (n=2). (B) Calcium flux of sorted Igh^g2A10^ cells incubated with Indo-1 dye and stimulated with CSP27, CSP9 or OVA-HEL, and the Ca2^+^ flux measured; near the end of the acquisition Ionomycin was added as a positive control; data shows the mean ± SD of 3 experimental replicates with summary data. (C) Summary data from B analysed via pairwise t-test, mean ± SD shown. (D) Total IgG response to NP measured via NP(14)BSA ELISA. (E) Representative flow cytometry plots showing gating of total IgD^-^ and GC B cells specific for NP or the CSP_Repeat_; values are percentages. (F) Absolute numbers of NP probe^+^ and CSP_Repeat_ tetramer^+^ IgD^-^ B cells. (G) Absolute numbers of NP probe^+^ and CSP_Repeat_ tetramer^+^ GC B cells. (H) Total IgG response to CSP_Repeat_ measured via (NANP)_9_ ELISA. Data for panels D-H are represented as mean ± SD pooled from two independent experiments (n≥4 mice/group/experiment); these data were analyzed via 2-way ANOVA, with experiment and mouse included in the model as fixed factors. ANOVA p values are listed below or adjacent to each graph. Pairwise comparisons were performed using a Tukey post-test and significant values are represented as symbols; * p<0.05, ** p<0.01, *** p<0.001.

To test whether the reduction in BCR signalling by CSP9 corresponded to a reduction in immunodominance of the CSP_Repeat_ we compared responses to CSP27-NP2 and NP-haptenated CSP9 (CSP9-NP2). CSP9-NP2 had significantly elevated NP-specific IgG compared to NP2-CSP27, particularly on days 14 and 21 post-immunisation (Fig. 3D). Analysis of the cellular NP and CSP_Repeat_ responses via flow cytometry at day 21 revealed that truncation of the CSP_Repeat_ not only reduced the number of cells responding to the CSP_Repeat_, but also allowed for a significant increase in the number of NP-specific total B cells and GC cells supporting the increase in antibody titres (Fig. 3E-G). Moreover, the CSP_Repeat_ IgG titres were significantly decreased for NP2-CSP9 immunised mice compared to NP2-CSP27 (Fig. 3H). Overall, this data supports the importance of valency in determining immunodominance as, decreasing the repeat length shifted the response away from the CSP_Repeat_ and towards NP.

We next performed the converse experiment and compared responses to CSP27, CSP27-NP2, CSP27-NP6 and CSP27-NP10. In these conditions the length of the CSP_Repeat_ antigen is fixed but the valency of NP varies. In agreement with the previous finding, increasing the NP:CSP ratio not only increased the response to NP (Fig S5A) but also decreased the level of antibodies to the CSP_Repeat_ (Fig S5B). Increasing the NP:CSP-ratio resulted in a switch in the immunodominance hierarchy; in mice immunized with CSP27-NP2, the CSP_Repeat_ response was immunodominant, while the NP response dominated the CSP_Repeat_ response upon immunization with CSP27-NP10 (Fig S5C-E). Collectively these data indicate a powerful role for antigen valency in drriving the immunodominance of repeating antigens.

### Repeat-truncated PfCSP molecules induce diverse antibody responses that protect against parasite challenge

One prediction of our data is that immunization CSP9 will induce stronger responses to the CSP_Nterm_ and CSP_Cterm_ compared to CSP27. Moreover, if these non-CSP_Repeat_ responses have anti-parasitic effect then immunization with CSP9 should be more protective than immunization with CSP27. Accordingly, we immunized mice 3 times at 5-week intervals with CSP9 and CSP27 formulated in alum before challenging mice with Pb-PfSPZ via mosquito bites 5 weeks after the final immunization (Fig. 4A). We also included an additional group which were immunized with CSP9_NVDP_ - a recombinant protein which also had a 9-mer repeat but included 3 NVDP repeats (Fig. S1) - as it has been suggested that (NANPNVDP)_n_ binding antibodies might confer superior protection to pure (NANP)_n_ binding antibodies (13).

**Fig. 4.**
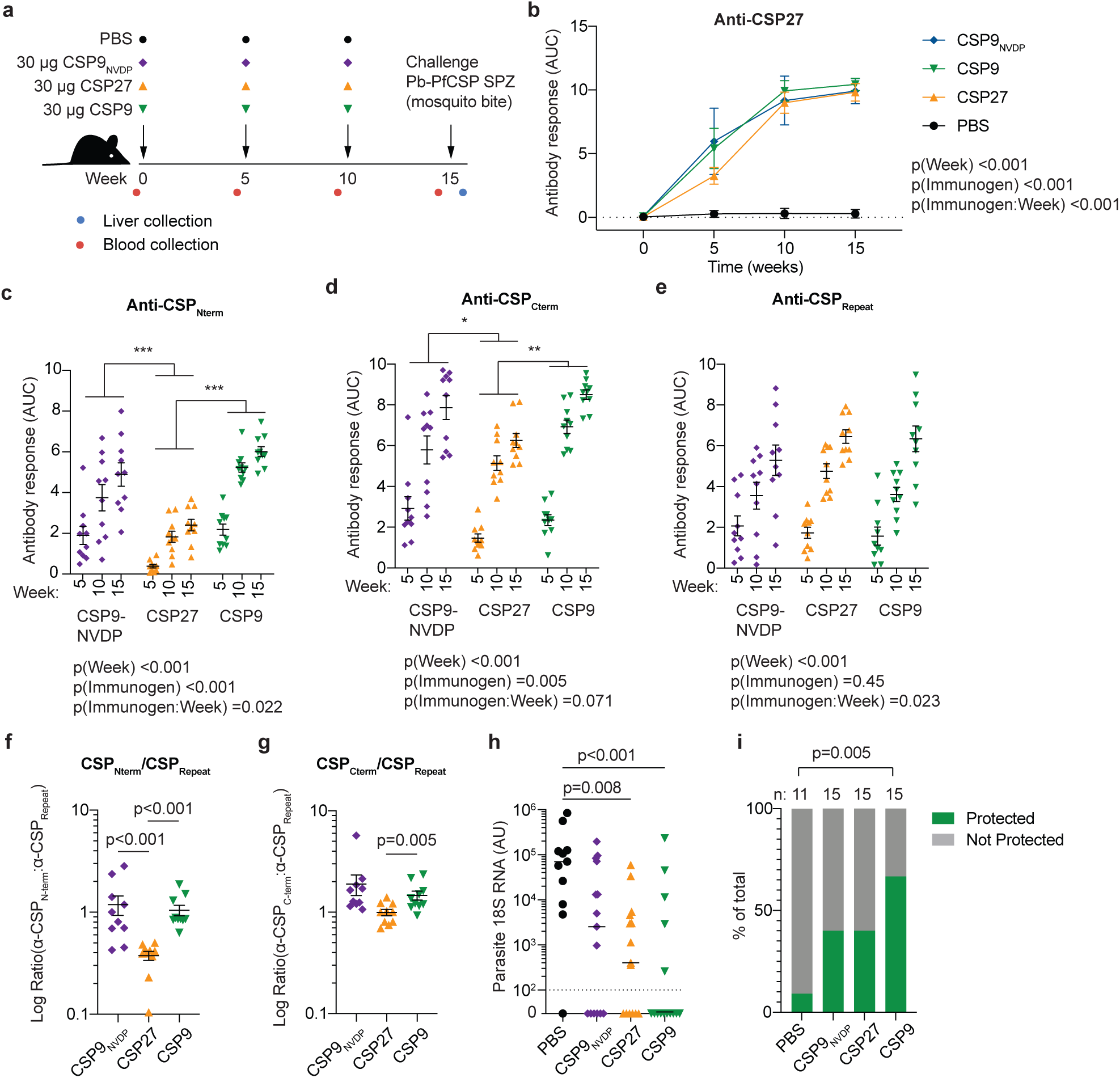
Immunization with CSP9 induces more diverse antibody responses and confers protection against sporozoite challenge. C57BL/6 mice were immunized three times at 5 weekly intervals with 30 µg CSP9, CSP9_NVDP_, CSP27 or alum only control, mice were challenged via mosquito bite with Pb-PfSPZ sporozoites and parasite burden measured by RT-PCR; blood was drawn prior to each immunization and challenge for analysis of the antibody response. (A) Schematic of the experiment. (B) Overall IgG responses to CSP27. (C) Overall IgG responses to CSP_Nterm_. (D) Overall IgG responses to CSP_Repeat_. (E) Overall IgG responses to CSP_Cterm_. Data in B-E was from experiments with 5 mice/experiment/group, analysed via 2-way ANOVA with experiment and mouse as blocking factors, ANOVA p values are listed below or adjacent to each graph; pairwise comparisons between groups (averaged over time) were performed using a Tukey post-test and significant values are represented as symbols; * p<0.05, ** p<0.01, *** p<0.001. (F) Ratio of the IgG response to CSP_Nterm_:CSP_Repeat_ at week 15 for the different immunized groups. (G) Ratio of the IgG response to CSP_Cterm_:CSP_Repeat_ at week 15 for the different immunized groups. Data for panels F and G were analyzed via one-way ANOVA with with experiment and mouse as blocking factors. (H) Parasite 18S RNA in the livers of mice 42 hours post challenge; data pooled from 3 experiments with 3-5 mice/experiment/group and analysed via Kruskal-Wallis test with Dunn’s multiple comparisons test. (I) Proportion of mice from (H) with no detectable parasite RNA in each group, pairwise comparisons were made via Fisher’s exact test.

Overall, IgG responses to *Pf*CSP (as measured via ELISA on CSP27 coated plates) were similar in magnitude across all immunized groups (Fig. 4B). Because differences between immunized mice and control mice are large (as are differences between pre-immune and post-immune sera), but not of particular interest, this group was removed from subsequent analysis as was the pre-immune timepoint to avoid skewing the statistical models. Examination of the specificity of the IgG responses at the domain level revealed significant differences in responses to the different immunogens: CSP9 and CSP9_NVDP_ induced significantly stronger IgG responses to CSP_Nterm_ and CSP_Cterm_ compared to CSP27 while responses to the repeat were similar (Figure 4C and D). We also performed more specific ELISAs with peptides corresponding to the regions bound by 5D5 and the CSP_Nterm_/CSP_Repeat_ junction (Fig. S6A). We found that the truncated CSP9 and CSP9_NVDP_ induced stronger responses to the 5D5 peptide which lies within the CSP_Nterm_ (Fig S6B) but that responses to the junction peptide were similar across all immunogens (Fig. S6C). Surprisingly, while CSP_Repeat_ specific antibody responses were initially lower in CSP9 immunized mice compared to CSP27 immunized mice, the repeat specific responses were similar after the third dose (Fig. 4E). Nonetheless, the ratio of antibody titres to the CSP_Nterm_ and CSP_Cterm_ domains to the CSPRepeat domains were significantly higher in the CSP9 immunized mice compared to CSP27 immunized mice at the time of challenge (Fig 4F and G).

Forty-two hours after mosquito bite challenge, livers were excised from the mice and the parasite burden measured by RT-PCR. Both CSP9 and CSP27 induced significant reductions in the mean parasite burden in the liver, whilst CSP9_NVDP_ did not (p = 0.069; Fig. 4H). Overall 10/11 (91%) control mice had detectable parasite 18SRNA, compared to only 5/15 (33%) CSP9 immunized mice which was statistically significant (p=0.005 by Fisher’s exact test; Fig. 4I). In both CSP27 and CSP9_NVDP_, 9/15 (60%) mice had detectable parasite RNA, which was not significantly different the control group (p=0.17 by Fisher’s exact test; Fig. 4I). The failure of CSP9_NVDP_ to induce better protection than the CSP9 was surprising to us, but did not reflect a failure to induce NVDP binding antibodies as these antibodies were significantly higher in both CSP27 (which includes NVDP repeats) and CSP9_NVDP_ immunized mice than in the CSP9 immunized mice (Fig. S6D and E). Overall, these data indicate that strategies which reduce the immunodominance of the CSP repeat can induce more diverse antibody responses and confer superior protection to vaccination strategies which utilise full length or near full length CSP.

## Discussion

Here we show that the primary driver of the immunodominance of the CSP_Repeat_ over other domains within CSP is the avidity of the binding between long repeats and BCRs on the surface of antigen specific B cells. Reducing the valency of the CSP_Repeat_ allows the development of stronger B cell and antibody responses to other epitopes. To demonstrate the importance of this observation for vaccination we used truncated CSP molecules to immunize against malaria in a pre-clinical model. We found that mice immunized with truncated CSP developed stronger responses to the CSP_Nterm_ and CSP_Cterm_ domains, both of which have been associated with protective antibody responses. Moreover, mice immunized with truncated CSP molecules were protected against live parasite challenge, and the magnitude of this protection was greater than in mice immunized with nearly full length CSP, though this difference did not reach statistical significance.

Our findings shed light on the key drivers of B cell immunodominance. There have been comparatively few studies on B cell immunodominance, but T cell immunodominance studies have highlighted critical roles for TCR-peptide MHC affinity and the number of naïve precursors specific for a particular antigen (reviewed in (32)). A common strategy for enhancing antibody responses is to generate polyvalent antigens, such as virus-like particles, to enhance immunity (33). Accordingly, we investigated how the multivalent nature of the CSP_Repeat_ affects the response to this epitope. Our finding that reducing the length of the CSP repeat allows responses to other antigens to develop suggests that not only does the long repeat drive a large response to this antigen itself, but also allows it to supress other responses. The discovery of this “immunodomination” effect provides direct support for the decoy hypothesis, though the exact mechanism for this effect is unclear. Our observation that CSP27 drives stronger BCR signalling than CSP9 suggests that this antigen may drive stronger expansion of the B cell response at the outset. An alternative – non-mutually exclusive hypothesis - is that is that CSP molecules carrying long repeats may be readily taken up by CSP_Repeat_ specific B cells, allowing these B cells to outcompete CSP_Nterm_ and CSP_Cterm_ specific B cells for T cell help (34, 35). In agreement with this, it has been previously shown that competition for restricted T cell help can supress responses to rare or subdominant epitopes (36).

We also investigated the roles of the number of antigen-specific precursors in determining the magnitude of the resulting immune response. The number of naïve precursors specific for a particular epitope was identified as a possible driver of CD4^+^ and CD8^+^ T cell immunodominance in mice in studies with peptide immunization or vesicular stomatitis virus infection (37, 38). However, other studies in humans (with HIV and hepatitis B virus) and mice (with influenza A virus) have not reported as strong a relationship as initially reported (39-41), perhaps due to compensation by other factors such as the affinity of the T cell receptor-MHC peptide interaction (32). In agreement with a previous study on the relationship between B cell precursor number and the subsequent GC response (22), we found that artificially increasing the number of precursors by transferring knock-in germline B specific for the CSP_Repeat_ cells did increase the magnitude of the GC response to this epitope. However, surprisingly, this enhanced response did not occur at the expense of responses to other epitopes and did not result in a significantly enhanced antibody response. One reason for the lack of immunodomination after the transfer of large numbers of CSP_Repeat_ specific B cells may be that precursor B cells for the competing antigen in this system (NP) are in excess. In agreement with this, NP specific B cells are reported to be present at high frequency (∼1/4000) in C57BL/6 mice (42). Thus, it will thus be important to determine if the same effect is seen when the number of precursors for the subdominant epitope is limiting.

Interest in developing “universal” vaccines for viruses such as influenza and HIV has led to increased focus on manipulating immunodominance hierarchies. This is because responses to conserved influenza epitopes (such as the HA stem) or conserved targets of bnAbs are subdominant. For HIV there is strong interest in developing multivalent immunogens targeting rare bnAb germline precursors to enhance the frequency of these cells (43, 44). For influenza, a possible vaccine strategy is to develop HA immunogens lacking the immunodominant, but variable head domain (23). One approach that might be possible, based upon studies in model systems would be to delete B cells specific for non-protective epitopes by injecting pure epitopes not linked to a T cell epitope (45). For CSP based malaria vaccines it is probably not desirable to remove the responses to the repeat altogether as antibodies targeting this domain are clearly protective, however a rebalancing of the immune response may be desirable. The current RTS,S vaccine may benefit from having a truncated repeat (of only 18 NANP moieties), though – critically - it lacks the CSP_Nterm_ domain and does not have any NVDP repeats (46). Perhaps surprisingly, we did not find that mice which mounted stronger responses to the NANPNVDP repeats were better protected than those which had responses more focussed on pure NANP repeats. This is despite the fact that mAbs that are strongly cross-reactive between both repeat motifs have been shown to protect against *Plasmodium* infection (13).

Collectively, our results provide insights into the factors that drive the immunodominance of different B cell epitopes; notably, that repeat epitopes can induce larger responses at the expense of subdominant epitopes. Our results suggest that truncated CSP molecules carrying all domains of the protein, may be a promising approach to the generation of a next generation CSP based vaccine.

## Materials and Methods

Full details of materials and methods used in this study are given in the supplementary information.

### Ethics statement

All animal procedures were approved by the Animal Experimentation Ethics Committee of the Australian National University (Protocol numbers: A2016/17; 2019/36). All research involving animals was conducted in accordance with the National Health and Medical Research Council’s Australian Code for the Care and Use of Animals for Scientific Purposes and the Australian Capital Territory Animal Welfare Act 1992.

### Mice and Parasites

C57BL/6 mice, B1-8 mice (31) and Ighg2A10 (McNamara et al. submitted) were bred in-house at the Australian National University. Mice were immunized IV with 5 x 10^4^ irradiated (15kRad) Pb-PfSPZ (29) dissected by hand from the salivary glands of *Anopheles stephensi* mosquitoes.

### Proteins and Immunizations

For immunization with CSP27, CSP9 or CSP9NVDP, or CSP27-NP conjugates 30 µg protein (or as described in the relevant figure legend) was emulsified in Imject™ Alum according to the manufacturer’s instructions (ThermoFisher Scientific) and delivered intra-peritoneally.

### Flow Cytometry

Single cell preparations of lymphocytes were isolated from the spleen of immunized mice and were stained for flow cytometry or sorting by standard procedures. Cells were stained with antibodies as outlined in Table S1 and CSP domain specific tetramers conjugated to PE or APC. Tetramers were prepared in house by mixing biotinylated (NANP)_9_ peptide with streptavidin conjugated PE or APC (Invitrogen) in a 4:1 molar ratio. Flow-cytometric data was collected on a BD Fortessa flow cytometer (Becton Dickinson) and analyzed using FlowJo software (FlowJo).

### ELISA

Binding of 2A10 antibody variants was determined in solid phase ELISA. Briefly, Nunc Maxisorp Plates (Nunc-Nucleon) were coated overnight with 1ug/ml streptavidin followed by binding of biotinylated peptide for 1 hour. After blocking with 1% BSA, serial dilutions of the antibodies were incubated on the plates for 1 hour and after washing, incubated with HRP conjugated anti-IgG antibodies (KPL). For the analysis of sera from immunized mice data were expressed as the area under the curve (AUC).

### Surface plasmon resonance

Surface plasmon resonance saturation experiments were performed on a Biacore 8K instrument (GE Healthcare) at 25 °C using a Series S Sensor Chip NTA (GE Healthcare). CSP27 and CSP9 were immobilized on separate channels on the sensor chip surface as per the manufacturer’s recommendations. A saturating solution of mAb 2A10 was then passed over the chip for 400 s, using a flow rate of 30 μl/min, followed by a 400 s dissociation period. The binding stoichiometry (*n*, molar ratio of antibody to antigen in the complex under saturating concentrations of mAb 2A10) was estimated as described previously (47).

### Quantitation of parasite RNA

*P. berghei* 18S rRNA was quantified from the livers of mice 42 hours after challenge via the bites of Pb-PfSPZ infected *Anopheles stephensi* mosquitoes as described previously (48).

### Statistical analysis

Statistical analysis was performed in GraphPad Prism for simple analyses without blocking factors; all other analyses was performed in R (The R Foundation for Statistical Computing) with details of statistical tests in the relevant figure legends.

## Acknowledgments

We thank Michael Devoy, Harpreet Vohra and Catherine Gillespie of the Imaging and Cytometry Facility at the John Curtin School of Medical Research for assistance with flow cytometry and multi-photon microscopy. We also thank Theresa Neeman of the ANU statistical consulting unit for assistance with statistical analysis of the data. This work was funded by the Bill and Melinda Gates Foundation (OPP1151018) and the National Health and Medical Research Council (GNT1158404). Authors declare no competing interests; All data is available in the main text or the supplementary materials.

