## Supplementary Information for "Avid binding by B cells to the *Plasmodium* circumsporozoite protein repeat suppresses responses to protective subdominant epitopes"

This supplement contains:

Supplementary Materials and Methods

Table S1: Antibodies used for flow cytometry in this study

Supplementary References

Figures S1-S6

#### Supplementary Materials and Methods

*Lymphocyte isolation and flow cytometry*

Spleens were collected from mice after cervical dislocation, and single cell suspension was prepared after passing through a 70 µM cell strainer into FACS buffer. Cells were briefly blocked for 30 minutes using 1µg/mL Streptavidin and 10 µg/mL TruStain fcX antibody diluted in FACS buffer. An antibody master mix was prepared in FACS buffer to stain the surface antigens of cells as per standard protocol using antibodies and B cell tetramers (prepared in house) listed in Table S1 for 30 minutes in dark. Cells were then lyzed using ACK lysis buffer and washed twice before staining with 1% 7AAD as Live/dead dye. Samples were acquired with a BD Fortessa or X20 flow cytometer, and data was analysed using FlowJo software. The universal gating strategy is shown Figure S2, after which the further gating to study the B cells of interest is provided in the results.

*Measuring antigen-specific sera IgG titres using ELISA*

Blood was collected from mice via either tail vein or retro-orbital bleeds and left to clot overnight at 4 °C, and the following day sera was collected by spinning the blood at 2000 *g* for 15 minutes at 4 °C. Next, Maxisorp Nunc-Nucleon 96 flat bottom plates were coated with 1µg/ml streptavidin overnight at 4 °C in dark. The following day, plates were washed in wash buffer consisting 0.05 % tween20 in PBS, and were incubated for 1 hour with different biotinylated peptides listed in pertaining to diverse sections of CSP protein. The peptide sequences were as follows, $CSP_{Nterm}$: Biotin-Ahx-NKNNQGNGQGHNMPNDPNRNVDENANANSAVKNNNNEEPSDKHIKEYLNKIQNSSTEWSPCSVTCGNGIQVRI KPGSANKPKDELDYANDIEKKICKMEKCS; $CSP_{Repeat}$: Biotin-Ahx-NANPNANPNANPNANPNANPNANP NANPNANPNANP, and $CSP_{Cterm}$:

Biotin-Ahx- QEYQCYGSSSNTRVLNELNYDNAGTNLYNELEMNYYG KQENWYSLKKNSRSLGENDDGNNEDNEKLRKPKHKKL KQPADG, additional peptides also carrying a Biotin-Ahx linker are described in Figure S6. Wells were blocked for 1 hour before incubating with initial sera (dilution of 1/100 times, followed by 1 in 4 serial dilutions down the plate) for one hour. After washing, anti IgG detection antibody conjugated to HRP was diluted 1/2000 times and was incubated for one hour. The plates were washed, then developed with Peroxidase Substrate Kit for 15 minutes, and the reaction was stopped using 50 µL/ well stop solution, consisting of 10% SDS in PBS. Absorbance at 405 nm (A405) was measured using an Infinite PRO Tecan plate reader, and the data was expressed as area under the curve (AUC) calculated in Prism 7 from the log(dilution) on the x axis and the A405 on the y axis, fitting a sigmoidal curve.

*Conjugation of NP to CSP*

CSP27 or CSP9 was initially concentrated using an Amicon ultra-15 centrifugal filter unit (10 kD molecular weight cut-off, MWCO). Briefly, the filter was washed by spinning 10 mL of MQ at 4000 *g* for 20 minutes at 4 °C then discarding the flow through and any retained water. 2 mg of CSP27 or 1 mg CSP9 was added to the membrane and made up to 10 mL in 3% $NaHCO_3$, then spun at 4000 *g* for 60 minutes at 4 °C. The flow-through was discarded and the retained protein made up to 10 mL in 3% $NaHCO_3$, then respun at 4000 *g* for 60 minutes at 4 °C. The flow-through was discarded and the retained protein was made up to final volume of 1 mL per mg of original starting material in 3% NaHCO+. This was then transferred into a pre-soaked 3.5 kD MWCO dialysis tubing and dialysed overnight, then for 4 hours in 1 L 3% $NaHCO_3$ at 4 °C.

Next, NP-ϵ-Aminocaproyl-Osu (NP-CAP-Osu) was dissolved in DMF to a final concentration of 10 mg/mL. We determined empirically that a 2:1 ratio of NP:CSP in the final conjugated product required a 20:1 ratio of NP:CSP at the conjugation step. For other NP:CSP ratios (6:1 and 10:1) this 10-fold increase during conjugation was also used; i.e. 60:1 and 100:1, respectively. The dialysed CSP and required NP-CAP-Osu were combined in sterile 2 mL tubes, covered with foil to protect from light, then rotated for 4 hours at RT to conjugate. The conjugated products were then dialysed in 1 L 3% $NaHCO_3$ at 4 °C for 5 hours, overnight then 4 hours before dialysis in 1 L PBS for 4 hours and overnight. Conjugated products were stored at 4 °C with 0.1% sodium azide to prevent bacterial growth.

*Absorption spectroscopy for quantifying NP*

Since NP-CAP-Osu has an absorption maximum of 430nm with extinction coefficient of 4230, the concentration of NP was determined via NanoDrop. Briefly, 1.5 µL aliquots of CSP27-NP/ CSP9-NP were analysed on a NanoDrop ND-1000 Spectrophotometer to measure protein at 280 nm and NP dye at 430 nm. An average of two reads was used to calculate the concentration of NP.

*Sandwich ELISA for calculating the concentration of CSP, post NP-CSP conjugation*

The absorbance of NP influenced the absorbance of proteins at 280 nm, therefore NanoDrop was not an accurate method for quantification of protein concentration (CSP27/CSP9) in the conjugated products. Instead, the concentration of CSP27 or CSP9 was determined by sandwich ELISA, taking advantage of histidine-tags (His-tag) attached to both CSP27 and CSP9 (Figure S1). Briefly, 96-well Maxisorp Nunc-Nucleon plates were coated overnight with 1 µg/mL of anti His-tag antibody. The following day, plates were washed and blocked with 1% BSA for 1 hour. The NP-CSP conjugated products were initially diluted 1/100 in 1%

BSA along with a 1/100 dilution of a CSP27 or CSP9 standard with known concentration, and serially diluted 1 in 5 times down the plate. Next, a 1/2000 dilution of 2A10 primary antibody (1) was incubated for one hour, washed and incubated with secondary antibody (Anti IgG detection antibody conjugated to HRP) for one hour. After washing, the plates were developed with Peroxidase Substrate Kit for 15 minutes and read at 405 nm using a Tecan Infinite 200Pro plate reader. The reaction was stopped using 50 µL/ well stop solution, consisting of 10% SDS in PBS. The concentration of CSP27 or CSP9 in the conjugated products was interpolated from a sigmoidal standard curve of CSP27 or CSP9.

*Preparing B cell tetramers for detecting diverse CSP specific or NP specific B cell response*

To detect B cells to specific epitopes in CSP we used tetramers based on peptide probes described above for the $CSP_{Repeat}$, $CSP_{Nterm}$, or $CSP_{Cterm}$ region conjugated to phycoerythrin (PE) or allophycocyanin (APC). The tetramers were generated by mixing biotin-conjugated peptides with streptavidin-conjugated PE or streptavidin-conjugated APC in a 4:1 molar ratio. Briefly, 2.17 nM of peptide was made up to 50 µL in PBS. Then 8.68 nM of PE or APC were added in 4 equal aliquots every 15 minutes, incubating at room temperature (RT) in darkness between aliquots. Tetramers were stored at 4 °C in dark until use.

NP-specific B cells were detected using 4-hydroxy-5-iodo-3-nitrophenol (NIP) conjugated to PE or APC. Briefly, 1 mg of Native R-Phycoerythrin protein or Natural Allophycocyanin protein were transferred into pre-soaked 3.5 kD MWCO dialysis tubing and dialysed for 5 hours, overnight, then for 4 hours in 1 L 3% $NaHCO_3$ at 4 °C. NIP- $\epsilon$ -Aminocaproyl-Osu (NIP-CAP-Osu) was dissolved in DMF to a concentration of 10mg/mL. The NIP-CAP-Osu was added to the dialysed PE or APC at a ratio of 20 µg:1 mg and rotated at RT for 4 hours protected from light with aluminium foil. The conjugated NP-PE and NP-APC were then dialysed in 1 L 3% $NaHCO_3$ at 4 °C for 5 hours, overnight then 4 hours before dialysis in 1 L PBS for 4 hours and overnight. NP probes were stored at 4 °C in dark till use.

*Immunizations with NP-CSP conjugates to study immunodominance*

Immunisations were conducted with the following amounts of antigen: NP-CSP27 = 15 µg/mouse, NP-CSP9 = 11.93 µg/mouse. Negative control groups were immunized with vehicle (PBS in alum). Antigens were emulsified in alum (2:1 volumetric ratio antigen: alum) to a total volume of 150 µL per mouse. The resultant solution was vortexed slowly at RT for 30 minutes to ensure the immunisations were fully emulsified. Immunisations were delivered intraperitoneally (IP), 150 µL total delivered in 75 µL aliquots on each side of abdomen into C57BL/6 recipient mice.

*Immunization with truncated CSP constructs*

Three different constructs of CSP were used for immunization, they include CSP9NVDP, CSP9, and CSP27 (Figure S1). C57BL/6 recipient mice were randomly separated in four groups, three of them had 15 mice per group, and were immunized with one of the CSP constructs. The fourth group had 11 mice and was the negative control. Each mouse received 30 µg of a CSP construct. These were emulsified in alum to a 2:1 volumetric ratio of antigen: alum and a resultant solution of 200 µL, before vortexing at RT for 30 minutes to ensure complete emulsification. Mouse were then IP immunized with 200 µL, 100 µL on left and right side, respectively. The immunisation regimen consisted of one priming and two boaster doses, each separated by an interval of 5 weeks. One day before immunization, blood was collected from mice via tail vein or retro-orbital bleeds for assessing antibody response. 2 weeks after the mice received their final booster, they were challenged via mosquito bite.

*Challenge of mice via* Anopheles stephensi *bites*

Controlled malaria infection challenge was performed via bite of *Anopheles stephensi* mosquitoes infected with Pb-PfSPZ a *P. berghei* parasite strain that expresses *P. falciparum* CSP (2). Also, parasite intrinsically expresses either GFP or mCherry, thus the infected mosquitoes were visually identified under microscope, and at least 5/4 of these were sorted onto separate containers and topped up with another 5/6. These were fed with sucrose for the first 6 hours and then with water solution a day before the challenge. The following day, mice were anaesthetised and placed on top of the containers to allow the mosquitoes to blood feed for 30 minutes. Next, 42 hours post mosquito bite, mice were euthanised via cervical dislocation and liver was collected, washed twice in PBS, and homogenised in 4 mL Denatured Working Stock.

*Quantification of parasite 18S rRNA in the liver*

To extract parasite rRNA, 60 µL of 2 M Sodium Acetate was added to a 600 µL aliquot of homogenised liver and vortexed to mix. Then 750 µL Acid Phenol:Chloroform was added and vortexed before incubating on ice for 15 minutes. The samples were spun at 15000 *g* for 20 minutes at 4 °C and the upper aqueous phase transferred into a clean 1.5 mL tube. The RNA was precipitated via addition of 400 µL isopropanol, vortexing then incubating at -20 °C for 1 hour. The RNA was pelleted at 15000 *g* for 20 minutes at 4 °C then the pellet washed twice with 1 mL cold 70% ethanol (EtOH) then dried for 10 minutes before resuspension in ultra-pure water. RNA concentration was measured via Nanodrop, and was diluted to make 100 µL aliquots at 50 ng/µL and stored at 4 °C.

cDNA was synthesised from the RNA using iScript cDNA Synthesis Kit according to the manufacturer's protocol. Briefly, each sample was run in a 20 µL reaction in individual dome-capped PCR tubes containing 10 µL ultra-pure water, 4 µL 5 x iScript Reaction Mix, 1 µL iScript Reverse Transcriptase and 5 µL RNA (50 ng/µL). The samples were run on an Eppendorf ProS Mastercycler at 25 °C for 5 minutes, 42 °C for 30 minutes, 85 °C for 5 minutes then hold at 4 °C.

RT qPCR was run using Power SYBR Green PCR Master Mix. Briefly, a PCR mastermix was made for all samples plus 2 no template controls (NTC) and 5 standards (STD) with the following volume per one reaction: 4.4 µL ultra-pure water, 5 µL Power SYBR Green PCR Master Mix, 0.05 µL *P. berghei* forward primer and 0.05 µL *P. berghei* reverse primer. 38 µL aliquots of mastermix were transferred into 1.5 mL tubes for each condition; 2 NTC, 5 STD and x cDNA samples. For DNA templates for *P. berghei* 18S qPCR 2 µL of the following was added for each condition; NTC – ultra-pure water, STD – plasmid standards ($10^7$, $10^6$, $10^5$, $10^4$ and $10^3$), samples – cDNA. The PCR was plated in triplicates of 10 µL/well in a MicroAmp 384 well reaction plate. The plate was sealed with MicroAmp optical adhesive film and spun at 500 *g* for 15 seconds to ensure all samples were at the bottom of each well. The qPCR was run on a 7900HT Fast Real-Time PCR System using the following conditions; 50 °C for 2 minutes, 95 °C for 10 minutes then 40 cycles of 95 °C for 15 seconds and 60 °C for 1 minute followed by 95 °C for 15 seconds and 60 °C for 15 seconds. The above qPCR reaction was repeated for glyceraldehyde 3-phosphate dehydrogenase (GapDH) using GapDH primers. However, the standards consisted instead of a pool of 2 µL cDNA from each sample that was serially diluted for the following concentrations; 1.0, 0.5, 0.25, 0.125, 0.0625. qPCR data was read using sDS2.4 software, *P. berghei* 18S was normalised to the GapDH reference gene before further calculating means in Prism 7.

*Calcium flux measurement in repeat specific B cells*

$CSP_{Repeat}$-specific B cells were FACS purified from $Igh^{g2A10}$ knock-in mouse splenocytes (McNamara *et al.* submitted), and were cultured in complete RPMI for 16 hours. Then the cells were labelled in RPMI media containing 2 μM/ml Indo-1 and 7AAD for 20 min at 37 °C. Following2 washes, the signal at BUV395 channel (Indo-1 bound) and BUV496 channel (Indo-1 free) was collected for 60 s to define baseline $Ca^{2+}$ levels as the ratio of Indo-1 (bound/free). B cells were then stimulated for additional 360 s with either 0.5 μM/ml CSP or 10 μg/ml OVA-HEL. 1 μg/ml ionomycin was used as positive control. Data were collected in a FACS Fortessa instrument (BD) and analyzed using the Kinetics tool in FlowJo software (Tree Star). To analyzed the result, the baseline $Ca^{2+}$ level was defined by the mean value of Indo-1 (bound/free) within 0-60 s. The calcium influx was then calculated by the mean values of every 30 s divided by the baseline value, and the plot was made by the calcium influx versus the mean value of this time frame.

*Surface Plasmon Resonance*

Surface plasmon resonance saturation experiments were performed on a Biacore 8K instrument (GE Healthcare) at 25 °C using a Series S Sensor Chip NTA (GE Healthcare) and SPR running buffer (10 mM HEPES, 150 mM NaCl, 50 μM EDTA, 0.05 % v/v Tween 20, pH 7.4). Solutions of $His_6$-tagged CSP27 and CSP9 were prepared in SPR running buffer at concentrations of 0.1 μg/ml and 0.4 μg/ml, respectively. $His_6$-tagged CSP27 and CSP9 were immobilized on separate channels on the sensor chip surface as per the manufacturer's recommendations: a pre-conditioned chip was first activated with 500 μM $NiCl_2$, and a solution of the $His_6$-tagged ligand was subsequently passed over the chip using a flow rate of 5 μl/min for 120 s. This yielded approximately 150 RU and 50 RU of immobilized CSP27 and CSP9, respectively. A saturating solution of mAb 2A10 (2 μM in SPR running buffer) was then passed over the chip for 400 s using a flow rate of 30 μl/min, followed by a 400 s dissociation period. The increase in response units (RU) corresponding to ligand immobilization ($RU_{lig}$) and analyte binding ($RU_{analyte}$) in the reference-subtracted (reference = blank surface) sensorgrams was measured, and the binding stoichiometry ($n$, molar ratio of antibody to antigen in the complex under saturating concentrations of mAb 2A10) was estimated using Eq. (1) as previously described (3), using molecular weights (MW) calculated using ProtParam (4):CSP27 (35.4 kDa), CSP9 (28.2 kDa) and mAb 2A10 (145.9 kDa).

$$n = \frac{RU_{analyte}}{RU_{ligand}} \times \frac{MW_{ligand}}{MW_{analyte}} \qquad (1)$$

All buffers were filtered and degassed prior to use. Following each cycle, the chip was completely regenerated using sequential washes of 500 mM imidazole, 350 mM EDTA (pH 8.5) and 100 mM NaOH. Each experiment was performed in duplicate (n=2), on separate channels on the SPR chip.

**Table S1: Antibodies used for flow cytometry in this study**

| Antibody | Conjugate | Clone | Source | Catalogue | Conc | Dilution |
|---|---|---|---|---|---|---|
| TruStain fcX antibody: Rat anti-mouse CD16/32 | - | 93 | Biolegend | 101320 | 0.5 mg/mL | 1/50 |
| Anti-mouse CD38 | APC A700 | 90 | eBioscience | 56-0381-82 | 0.2 mg/mL | 1/200 |
| Anti-mouse IgM | APC Cy7 | II/41 | Invitrogen | 47-5790-82 | 0.2 mg/mL | 1/200 |
| Anti-mouse CD45.2 | APC Cy7 | A20 | Biolegend | 110716 | 0.2 mg/mL | 1/200 |
| Anti-mouse CD19 | BUV395 | 1D3 | BD Horizon | 563557 | 0.2 mg/mL | 1/200 |
| Anti-mouse IgD | BV605 | 11-26c.2a | Biolegend | 405727 | 0.2 mg/mL | 1/400 |
| Anti-mouse B220 | BV605 | RA3-6B2 | BioLegend | 103244 | 0.2 mg/mL | 1/200 |
| Anti-mouse CD45.1 | FITC | A20 | Biolegend | 110706 | 0.5 mg/mL | 1/200 |
| Anti-mouse CD45.2 | FITC | 104 | Invitrogen | 11-0454-82 | 0.5 mg/mL | 1/200 |
| Anti-mouse GL7 | Pacific Blue | GL-7 | Invitrogen | 48-5902-82 | 0.2 mg/mL | 1/100 |
| Anti-mouse CD45.2 | PE | 104 | Biolegend | 109807 | 0.2 mg/mL | 1/200 |
| Anti-mouse CD138 | PE Cy7 | 281-2 | Biolegend | 142514 | 0.2 mg/mL | 1/100 |
| Anti-mouse CD3 | PerCP Cy5.5 | 17A2 | Biolegend | 100218 | 0.2 mg/mL | 1/200 |
| Anti-mouse CD11b | PerCP Cy5.5 | M1/70 | Biolegend | 101228 | 0.2 mg/mL | 1/200 |
| Anti-mouse CD11c | PerCP Cy5.5 | N418 | Biolegend | 117328 | 0.2 mg/mL | 1/200 |
| Anti-mouse Ly-6G/Ly-6C (GR1) | PerCP Cy5.5 | RB6-8C5 | Biolegend | 108428 | 0.2 mg/mL | 1/200 |
| 7AAD Cell Viability Dye | PerCP Cy5.5 | - | Biolegend | 420404 | 50 □g/mL | 1% |

### Supplementary Figure 1

**a**

```
   >CSP 3D7 genome sequence
  1 MMRKLAILSV SSFLFVEALF QEYQCYGSSS NTRVLNELNY DNAGTNLYNE
 51 LEMNYYGKQE NWYSLKKNSR SLGENDDGNN EDNEKLRKPK HKKLKQPADG
101 NPDPNANPNV DPNANPNVDP NANPNVDPNA NPNANPNANP NANPNANPNA
151 NPNANPNANP NANPNANPNA NPNANPNANP NANPNANPNA NPNANPNVDP
201 NANPNANPNA NPNANPNANP NANPNANPNA NPNANPNANP NANPNANPNA
251 NPNANPNANP NANPNANPNA NPNKNNQGNG QGHNMPNDPN RNVDENANAN
301 SAVKNNNNEE PSDKHIKEYL NKIQNSLSTE WSPCSVTCGN GIQVRIKPGS
351 ANKPKDELDY ANDIEKKICK MEKCSSVFNV VNSSIGLIMV LSFLFLN
```

**b**

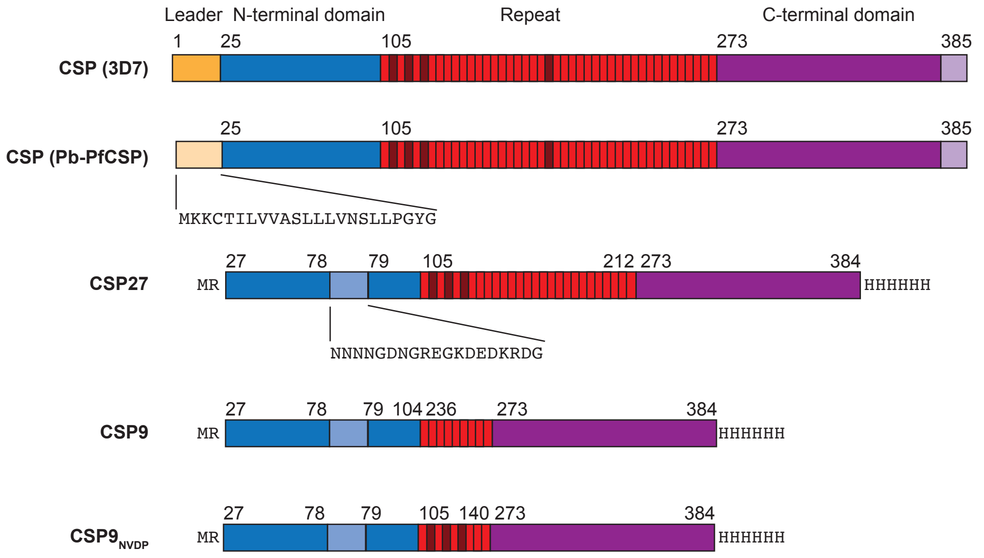


**Supplementary Figure 1: Circumsporozoite protein sequences and constructs used in this study**
(A) Sequence of the 3D7 circumsporozoite protein, different domains are labelled with different colours: leader sequence, orange; N-terminal domain, blue; repeat, NANP – light red, NVDP – dark red; C terminal domain, purple. (B) Schematics of the different constructs used in this study, numbers refer to the 3D7 sequence, insertions/substitutions are marked as text.

#### Supplementary Figure 2

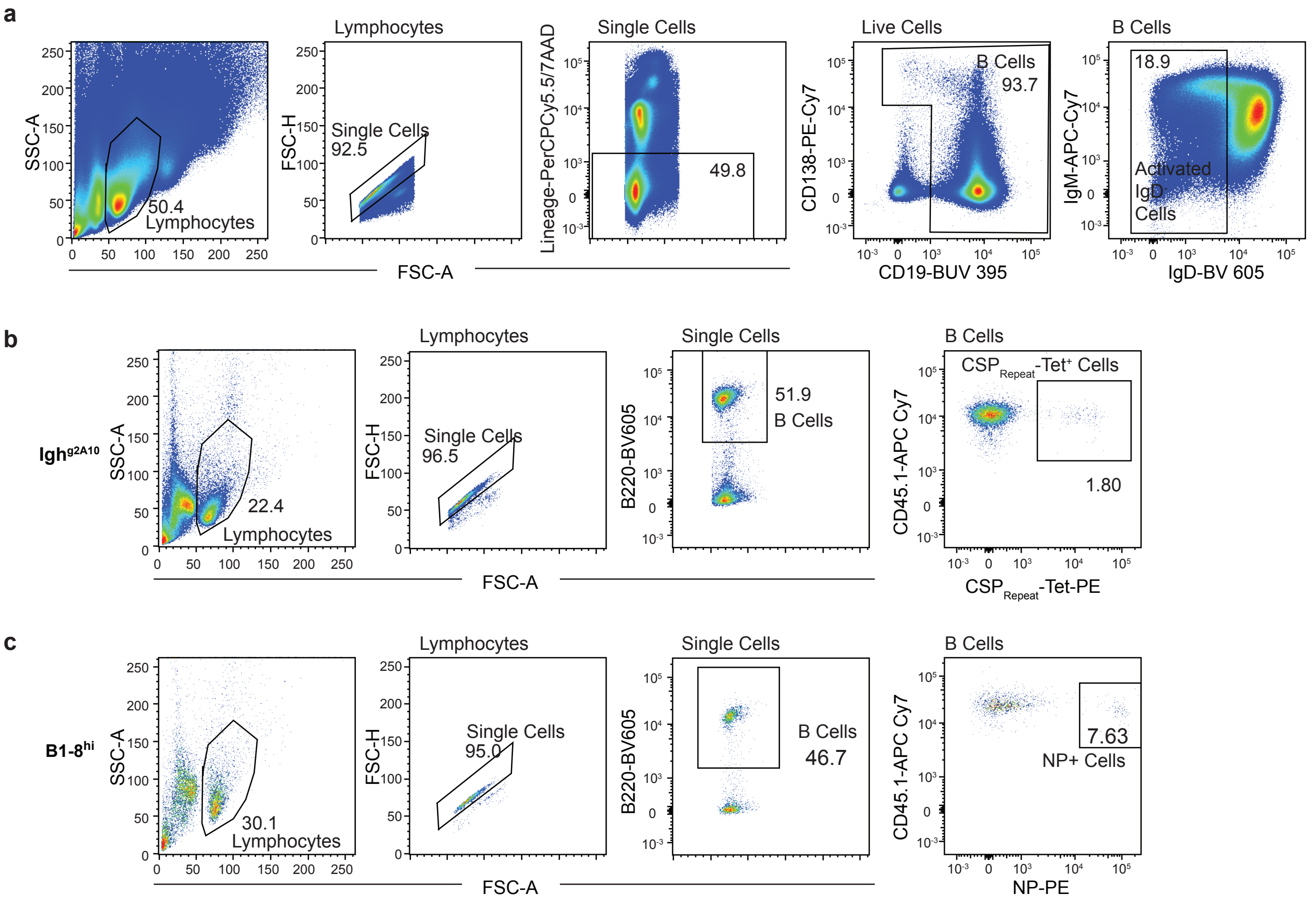


**Supplementary Figure 2: Gating strategies for flow cytometry** (A) General gating strategy applied to all flow cytometry data prior to gating antigen specific cells. Firstly, lymphocytes were gated based on their relative size and granularity (forward scatter area versus side scatter area). From the lymphocyte population, single cell events were gated on relative forward scatter area versus forward scatter height. From the single cell population, live cells were gated on PerCP Cy5.5 negative population (this is negative for 7AAD viability dye and also negative for lineage markers CD3, CD11b, CD11c and GR1). From the live cells, B cells and plasmablasts were gated as CD19$^{+}$ and/or CD138$^{+}$ cells. From the B cell population, activated cells were gated as IgD$^{-}$ cells. The activated cells were then further gated for antigen specificity. Plots of further gating are provided in results figures. (B) Gating strategy for quantifying Igh$^{g2A10}$ NANP tetramer+ B Cells. Lymphocytes were gated based on their relative size and granularity (forward scatter area versus side scatter area). From the lymphocyte population, single cell events were gated on relative forward scatter area versus forward scatter height. From the single cell population, B cells were gated as B220$^{+}$ cells. From the B cell population, CSP$_{Repeat}$-tet$^{+}$ cells were gated as cells double-positive for both the tetramer and congenic marker CD45.1. (C) Gating strategy for quantifying B1-8$^{Hi}$ NP+ B cells. The lymphocyte, single cell and B cell populations were gated as described in (A). From the B cell population, NP$^{+}$ B cells were gated as cells double-positive for both the NP marker and congenic marker CD45.1.

#### Supplementary Figure 3

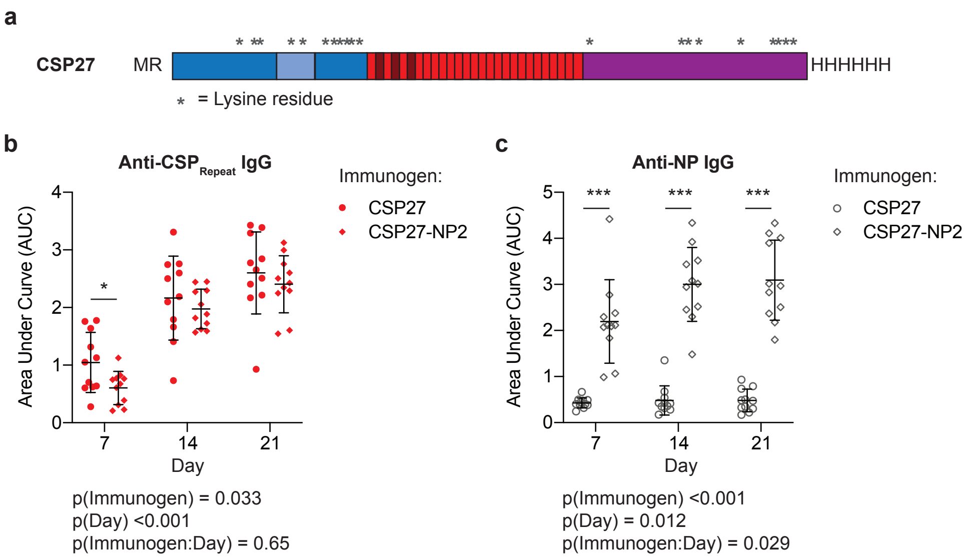


**Supplementary Figure 3: CSP27-NP2 elicits both CSPRepeat and NP specific IgG responses** C57BL/6 mice were immunised with CSP27-NP2 or CSP27 only control. Sera were taken 7, 14 and 21 days post-immunisation and the antibody response to each epitope measured via ELISA. (A) Schematic of CSP27 showing the locations of lysine residues, indicated by asterisks. (B) Total IgG response to $CSP_{Repeat}$ measured via $(NANP)_9$ ELISA. (C) Total IgG response to NP measured via NP(14)BSA ELISA. Data are represented as mean ± SD pooled from three independent experiments (n≥3 mice/group/experiment); ELISA data were analysed separately for each antigen in R Studio using a mixed linear model with Immunogen (CSP27-NP2/CSP27) and day as experimental factors, experiment as a fixed factor and mouse as a random factor. Two-way ANOVA p values are listed underneath each graph, pairwise comparisons were made via Tukey post test and are represented using symbols; * p<0.05, ** p<0.01, *** p<0.001.

#### Supplementary Figure 4

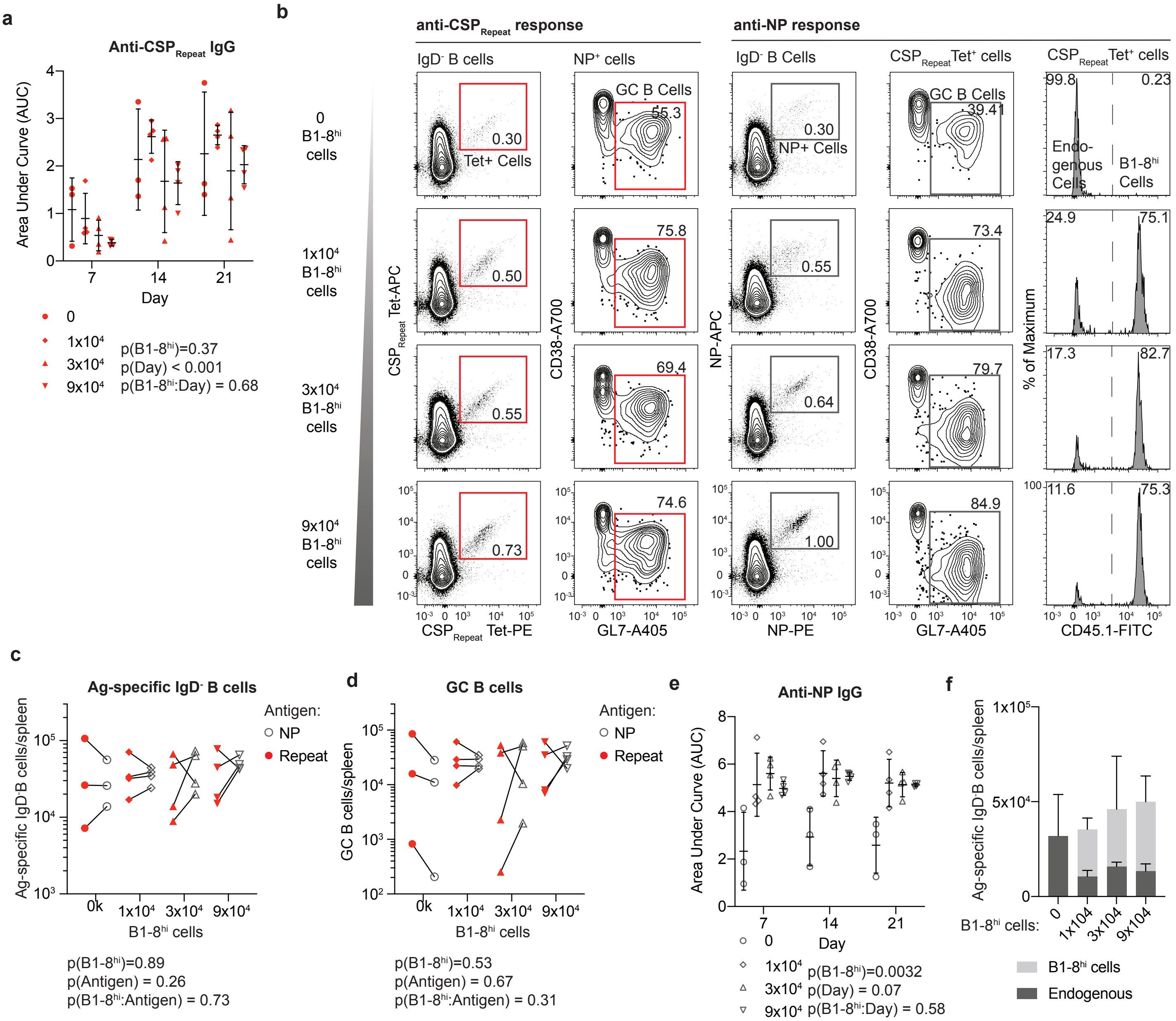


**Supplementary Figure 4: Increasing NP-specific precursor number does not alter immunodominance.** 0, $1\times10^4$, $3\times10^4$, or $9\times10^4$ of CD45.1 B1-8$^{hi}$ cells were adoptively transferred into C57Bl/6 mice followed by immunization with CSP27-NP2 in alum. Sera were taken on days 7, 14 and 21 and spleens analyzed 21 days post-immunization. (A) Total IgG response to CSP$_{Repeat}$ measured via (NANP)$_9$ ELISA. (B) Representative flow cytometry plots showing gating of total IgD$^-$ and GC B cells specific for NP or the CSP$_{Repeat}$; values are percentages (C) Absolute numbers of NP probe$^+$ and CSP$_{Repeat}$ tetramer$^+$ IgD$^-$ B cells. (D) Absolute numbers of NP probe$^+$ and CSP$_{Repeat}$ tetramer$^+$ GC B cells. (E) Total IgG response to NP measured via NP(14)BSA ELISA. (F) Absolute numbers of CSP$_{Repeat}$ tetramer$^+$ CD45.1$^+$ Igh$^{g2A10}$ and CD45.1$^-$ endogenous cells. Data are represented as mean ± SD from a single experiment (n=3 mice/group); all data were analyzed via 2-way ANOVA, with mouse included in the model as a fixed factor. ANOVA p values are listed below or adjacent to each graph. Pairwise comparisons were performed using a Tukey post-test though no pairwise differences were significant.

#### Supplementary Figure 5

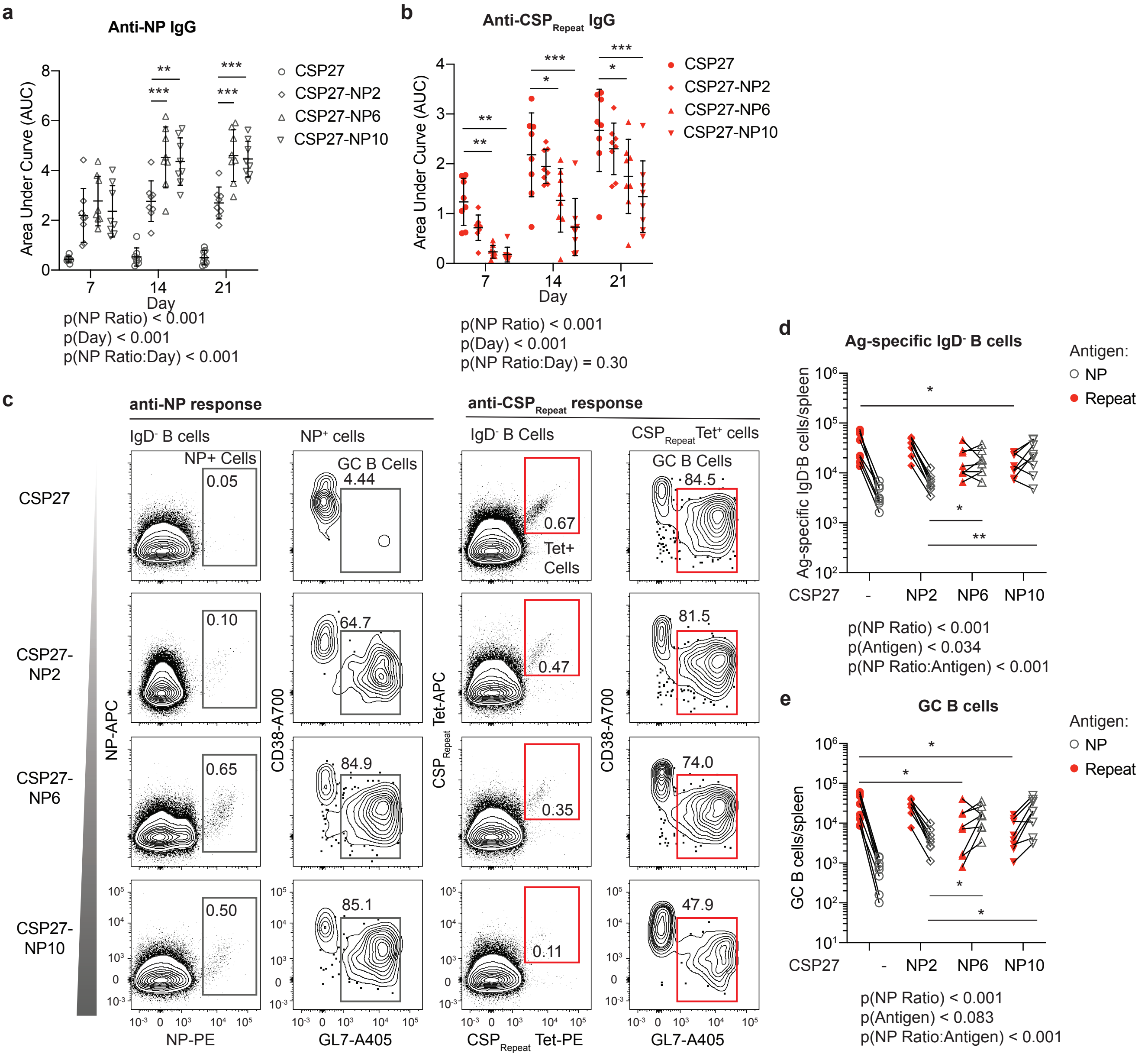


**Supplementary Figure 5: Increasing the level of NP conjugation to CSP alters the immunodominance hierarchy** C57BL/6 mice were immunized with either CSP27, CSP27-NP2, CSP27-NP6 or CSP27-NP10. Sera were taken on days 7, 14 and 21 and spleens analyzed 21 days post-immunization. (A) Total IgG response to NP measured via NP(14)BSA ELISA. (B) Total IgG response to $CSP_{Repeat}$ measured via $(NANP)_9$ ELISA. (C) Representative flow cytometry plots showing gating of total $IgD^-$ and GC B cells specific for NP or the $CSP_{Repeat}$; values are percentages (D) Absolute numbers of NP $probe^+$ and $CSP_{Repeat}$ $tetramer^+$ $IgD^-$ B cells. (E) Absolute numbers of NP $probe^+$ and $CSP_{Repeat}$ $tetramer^+$ GC B cells. Data are represented as mean ± SD pooled from two independent experiments (n=4 mice/group/experiment); these data were analyzed via 2-way ANOVA, with experiment and mouse included in the model as fixed factors. ANOVA p values are listed below or adjacent to each graph. Pairwise comparisons were performed using a Tukey post-test with significant values are represented as symbols; * p<0.05, ** p<0.01, *** p<0.001.

### Supplementary Figure 6

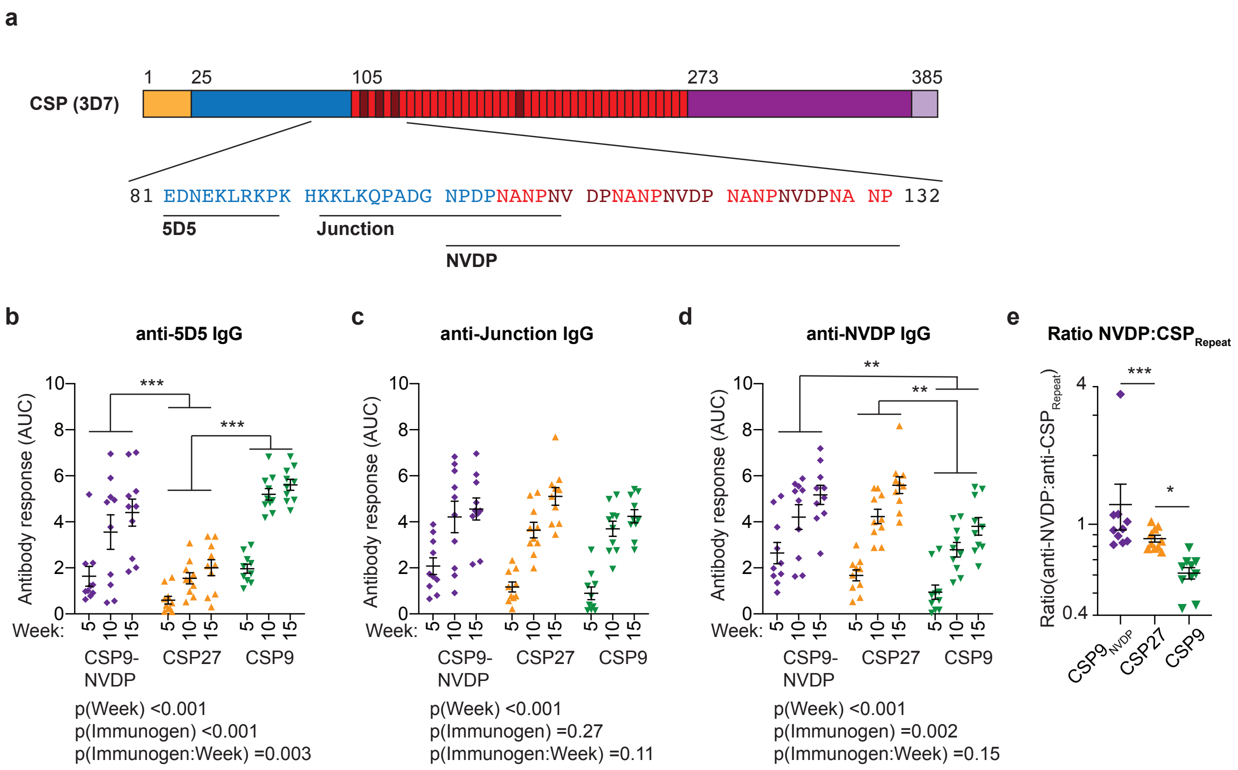


**Supplementary Figure 6: Additional analysis of mice immunized with truncated CSP molecules.** Sera from the mice described in Figure 4A were taken for additional ELISA analysis. (A) Sequence and location within the CSP molecule of additional peptides corresponding to potential targets of protective antibodies. (B) Overall IgG responses to the 5D5 epitope. (C) Overall IgG response to the junction between the CSPNterm and CSPRepeat. (D) Overall IgG response to a peptide corresponding to the first 32 amino acids of the 3D7 CSP repeat domain including multiple NVDP repeats. (E) Ratio of the week 15 response to the NVDP peptide and the standard $CSP_{Repeat}$ peptide between the different immunization groups. Data from panels B-D was analyzed from 2 experiments with 5 mice/experiment/group analyzed via 2-way ANOVA with experiment and mouse as blocking factors, ANOVA p values are listed below or adjacent to each graph; pairwise comparisons between groups (averaged over time) were performed using a Tukey post-test and significant values are represented as symbols; * $p<0.05$, ** $p<0.01$, *** $p<0.001$. Data from panel E was from 2 experiments with 5 mice/experiment/group analyzed via one-way ANOVA with experiment as a blocking factor.
